# Fragment-based computational design of antibodies targeting structured epitopes

**DOI:** 10.1101/2021.03.02.433360

**Authors:** Mauricio Aguilar Rangel, Alice Bedwell, Elisa Costanzi, Ross Taylor, Rosaria Russo, Gonçalo J. L. Bernardes, Stefano Ricagno, Judith Frydman, Michele Vendruscolo, Pietro Sormanni

## Abstract

De novo design methods hold the promise of reducing the time and cost of antibody discovery, while enabling the facile and precise targeting of predetermined epitopes. Here we describe a fragment-based method for the combinatorial design of antibody binding loops and their grafting onto antibody scaffolds. We designed and tested six single-domain antibodies targeting different epitopes on three antigens, including the receptor-binding domain of the SARS-CoV-2 spike protein. Biophysical characterisation showed that all designs are highly stable, and bind their intended targets with affinities in the nanomolar range without any *in vitro* affinity maturation. We further discuss how a high-resolution input antigen structure is not required, as our method yields similar predictions when the input is a crystal structure or a computer-generated model. This computational procedure, which readily runs on a laptop, provides a starting point for the rapid generation of lead antibodies binding to pre-selected epitopes.

**summary:** A combinatorial method can rapidly design nanobodies for predetermined epitopes, which bind with KDs in the nanomolar range.

---

Antibodies are key tools in biomedical research, and are increasingly employed to diagnose and treat a wide range of human diseases. Currently, there are over 130 antibodies s approved or undergoing regulatory review in the United States or European Union (*1*). Existing antibody discovery methods have been widely successful, but still have important limitations (*2*). Extensive laboratory screenings are required to isolate those antibodies binding to the intended target, which can be time consuming and costly. Some classes of hard targets remain, including some receptors and channels, proteins within highly homologous families, aggregation-prone peptides, and disease-related short-lived protein aggregates (*3, 4*). Most notably, it is often highly challenging to obtain antibodies targeting pre-selected epitopes. Screening procedures typically select for the tightest binders, which usually occur for immunodominant epitopes, thus disfavouring the discovery of antibodies with lower affinities but binding to functionally relevant sites (*5*). Furthermore, screening campaigns often yield antibodies with favourable binding affinity, but otherwise poor biophysical properties, such as stability, solubility, and production yield, which may hinder their development into effective reagents. Computational antibody design has the potential to overcome these limitations by drastically reducing time and costs of antibody discovery, and in principle allowing for a highly controlled parallel screening of multiple biophysical properties. Importantly, rational design inherently enables the targeting of specific epitopes.

Most available methods for the design of binding proteins rely at least in part on the minimisation of a calculated interaction free energy, through the sampling of the mutational and conformational space (*2, 6, 7*). The nature of these calculations, which are based on molecular modelling and classical force fields, and the challenges of achieving exhaustive sampling, make design simulations rather imprecise and highly resource intensive. For these reasons, the de novo design of antibody binding has generally met low success rates, and required recursive experimental screenings and large libraries (*5, 8*–*10*), which hamper its competitiveness with established laboratory-based technologies. Computational design of binding has been most successful in synergy with *in vitro* affinity maturation, and in particular when applied to mini-proteins (*11, 12*). The small size of these mini-proteins is amenable to the high-throughput gene synthesis required to experimentally screen designed candidates on a massive scale, and their rigidity reduces the need for accurate conformational sampling. However, antibody domains are considerably larger, and bind their target using complementarity determining regions (CDRs) located within hypervariable loops on the antibody surface, which are often extended and highly flexible.

Here, we describe a novel method to design antibody CDR loops targeting epitopes of known structures, or for which a structural model is available. Designed CDRs are then grafted onto antibody scaffolds, and further optimised computationally for solubility and conformational stability. Novel antibody-antigen interactions are designed by combining together protein fragments identified as interacting with each other within known protein structures.

## De novo CDR-design strategy

To overcome some of the limitations of molecular modelling mentioned above, in particular those associated with the approximations in accounting for interatomic interactions, we exploited the availability of large structural databases to implement a fragment-based procedure to design CDRs (paratope) complementary to a target epitope. To implement this idea, we compiled from the non-redundant Protein Data Bank (PDB) a database of CDR-like fragments and corresponding antigen-like regions, which we call *AbAg database*. CDR-like fragments are defined as linear motifs structurally compatible with an antibody CDR loop, which may be found in any protein structure in the PDB, and antigen-like regions are those found interacting with any CDR-like fragment in the structures analysed (see Supplementary Methods).

The structure of the input epitope is fragmented using two different strategies (**Fig. 1A**): (i) a linear fragmentation, which generates fragments of at least four consecutive residues, and (ii) a surface-patch fragmentation, which takes each residue and yields the closest *n*≥4 solvent-exposed residues in the three-dimensional structure of the epitope. Next, each epitope fragment is compared to the antigen-like regions to identify those with compatible backbone structure and similar sequence. More specifically, the search is carried out with the Master algorithm (*13*) and the comparison is based on the root-mean-square deviation (RMSD) of the full backbone as well as on sequence similarity (see Supplementary Methods). Therefore, a hit antigen-like region is similar to its query epitope fragment in both sequence and structure. In practice, the fragmentation is carried out starting from large fragments (i.e. from the full region defined as epitope), and moving to smaller ones for a minimum size of four residues. Most commonly, no hits are found for larger fragments, while many hits are typically found for smaller ones (*n ≤ 6*).

**Figure 1.**
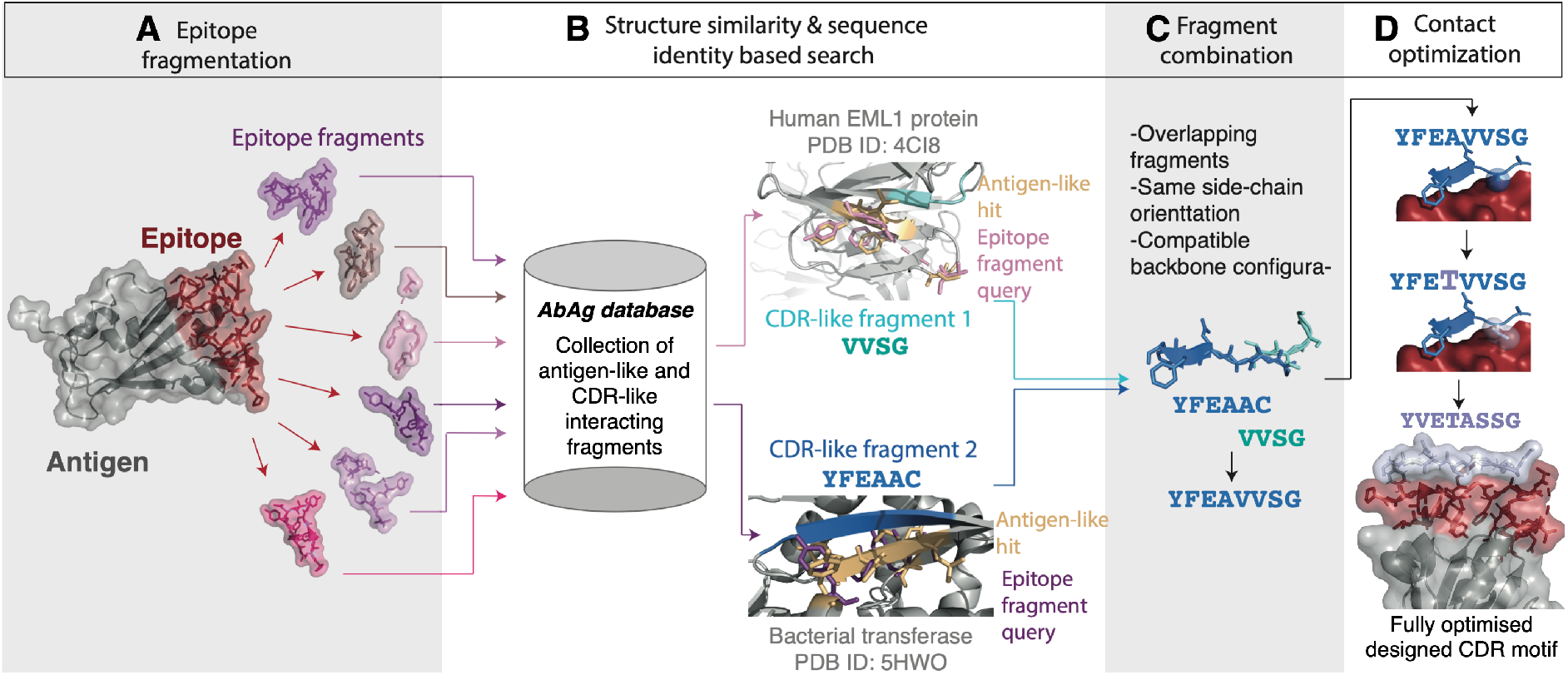
Workflow for the combinatorial structure-based CDR design strategy. (**A**) The antigen structure is shown in grey with the epitope of interest highlighted in red. At this step the epitope is fragmented into its structural fragments, both in a linear mode and in a surface-patch mode that yields also non-contiguous fragments (see text). Some example fragments are pointed by red arrows. (**B**) These fragments are used as queries for a structural search against a custom database of structures of antigen-like and CDR-like interacting fragments. Hits are selected based on structural and sequence similarity with the query epitope fragments, and two example hits are depicted: an epitope fragment (pink top example, purple lower example) matching antigen-like fragments (yellow) interacting with a CDR-like fragment (cyan top example, blue lower example). These two examples originate respectively from the structures of human EML1 protein and of a bacterial transferase, as antigen-like and CDR-like fragments may be found in any structure from the PDB. (**C**) When possible, identified CDR-like fragments are joined together. Here, the overlapping CDR-like fragments from B are merged as they meet the stated compatibility criteria. (**D**) The sequence of the designed CDR fragment resulting from the merging is optimized to increase the probability of favourable CDR-epitope contacts. The final fully optimised designed CDR motif can then be grafted onto suitable antibody frameworks. The example in this figure corresponds to the designed binding motif within the CDR3 of DesAb-RBD-C1 targeting the ACE2-binding-site on the RBD domain.

Because of the structure of the *AbAg database*, this procedure automatically yields those CDR-like fragments that interact with the identified antigen-like fragments (**Fig. 1B**). These CDR-like structures are then rotated to match the orientation of the epitope, by superimposing each antigen-like region, together with its interacting CDR-like fragments, to the matching part of the epitope (**Figs. 1** and **S1**). When possible, different CDR-like fragments whose backbones are partly overlapping and compatible with a single longer CDR loop are joined together to yield longer interacting motifs (**Fig. 1C**, see Supplementary Methods).

Some of the original interactions of each CDR-like fragment may be affected when this fragment is transferred onto the epitope, for instance if the sequence of the antigen-like region is not identical to the corresponding epitope sequence, or if the epitope sidechains are found in different conformations (**Fig. 1D**). Similarly, new interactions may arise when a CDR-like fragment forms contacts with parts of the epitope that were not matched onto its antigen-like region. To overcome potential issues arising from these suboptimal interactions, we implemented a side-chain optimisation procedure that seeks to maximize the number of favourable interactions between the CDR-like fragment ant the antigen. Briefly, for each CDR-like sidechain with interactions different, or additional, to those found in the original hit, a structural neighbourhood is defined by taking the backbone coordinates of all contacting residues (see Supplementary Methods). Such residues are then used as a query to interrogate the *AbAg database*, retrieving as hits those CDR-like sidechains that better match the native local environment of the epitope, therefore increasing the total number of favourable interactions to yield a fully optimised designed CDR motif (**Fig. 1D**, see Supplementary Methods).

Typically, multiple CDR motifs are designed in this way for a given input epitope, as multiple CDR-like fragments are usually identified as suitable starting points for the combination and the optimisation procedures. Therefore, all possible CDR-motif candidates generated for the input epitope are ranked according to the total number of favourable interactions, the number of interactions that could not be optimised, and a solubility score calculated with the CamSol method (*14*).

Top-ranking, fully optimised designed CDR motifs can then be grafted into an antibody scaffold (**Fig. 2**). Our pipeline can structurally match the generated motifs to either complete CDRs or entire antibody (specifically Fv regions) structures, which can result in longer CDR loops harbouring multiple motifs, or in multiple motifs being grafted in different CDR loops of the same Fv region (**Fig. 2A-C**, see Supplementary Methods). If needed, any new interactions between the grafted antibody scaffold and the antigen are optimised using the side-chain optimisation procedure described above. Furthermore, as an alternative to this structural matching, fully optimised designed CDR motifs can also be grafted directly into an antibody scaffold that is already known to be highly tolerant to loop replacements. In this work we tested experimentally both approaches (**Fig. 2D**).

**Figure 2.**
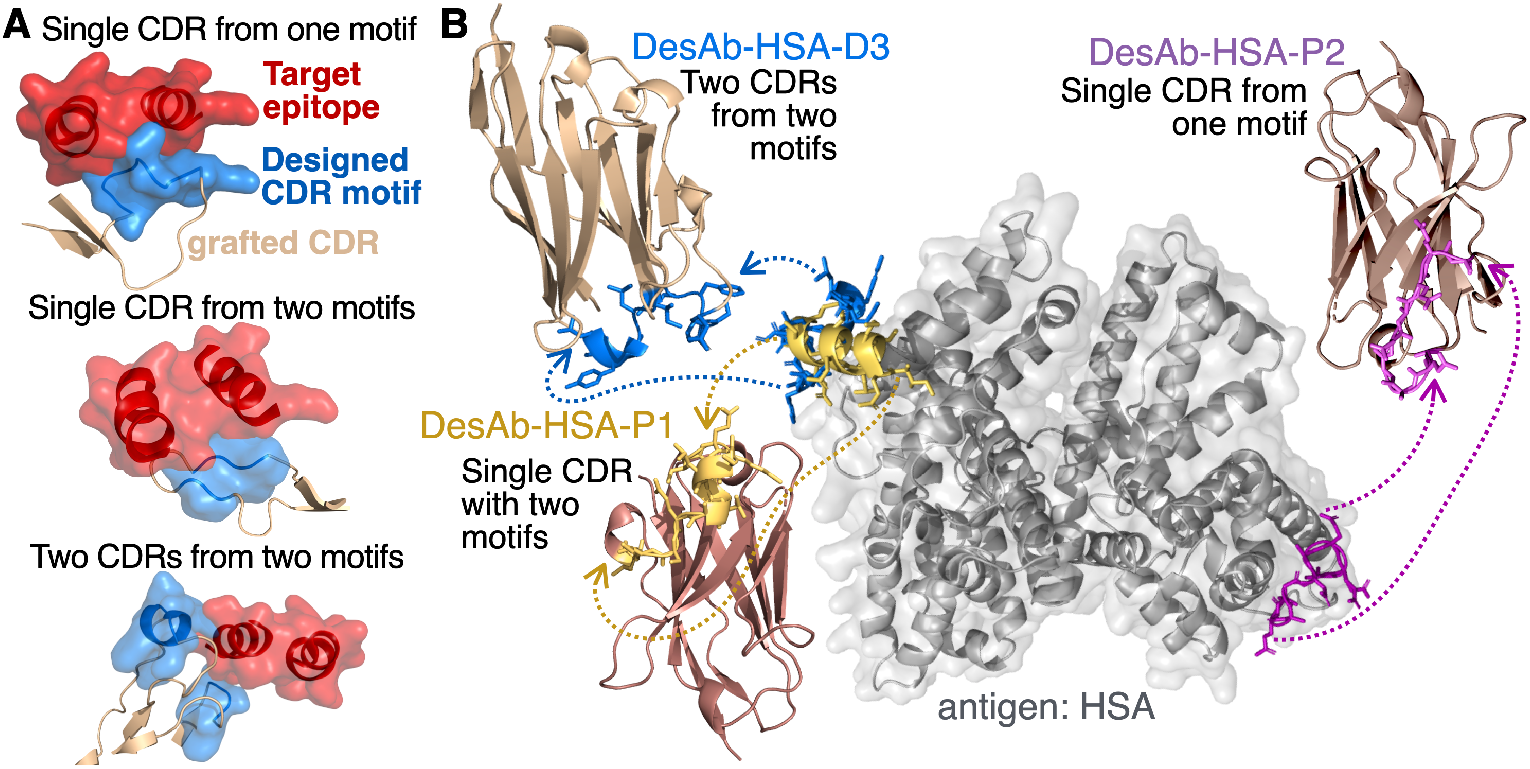
Grafting of designed CDR motifs onto antibody scaffolds. (**A-C**) Three examples of how designed CDR motifs can be grafted in different ways. The epitope is shown in red and the designed CDR motifs in light blue. These are grafted on to structurally matched CDR loops (light brown). In (**A**) a single motif is grafted in a loop, in (**B**) two motifs are grafted on the same loop, and in (**C**) two motifs are grafted in two different loops from the same Fv domain. Multiple CDR-like fragments are joined in a single motif when overlapping (like in Fig. 1C), or, if not overlapping, may still be grafted in the same CDR loop as shown in (**B**). (**D**) The structure of HSA is shown in grey and the designed CDR motifs selected for experimental validation are shown in blue, yellow, and purple docked onto their respective epitopes. Two fragments (blue) are grafted into separate CDRs (CDR1 and CDR3) of an antibody scaffold which they match structurally (PDB 4DKA). The resulting design is DesAb-HSA-D3 (Table 1). The yellow and purple motifs are instead grafted into the CDR3 of a scaffold resilient to CDR3 substitutions to yield respectively DesAb-HSA-P1 and DesAb-HSA-P2. The motif grafted onto DesAb-HSA-P1 comprises two fragments joined together as in Fig. 1C. DesAb structural models were obtained with the SAbPred webserver(*34*).

To validate our design strategy, we tested it experimentally on single-domain antibodies, because of their monomeric nature, ease of production in prokaryotic systems, and small size (*15*). Nonetheless, the computational design pipeline described here can readily be applied to other antibody fragments, including whole Fv regions, on which designed CDR motifs can be structurally matched and grafted in the same way on either heavy or light chain CDRs.

## Description of designs and biophysical characterization

We designed six single-domain antibodies for three different antigens by exploring two grafting strategies: the structural matching of designed CDR motifs, and the simpler grafting into stable scaffolds. Two designed single-domain antibodies target the SARS-CoV-2 spike protein receptor-binding domain (RBD), three human serum albumin (HSA), and one pancreatic bovine trypsin (**Table 1**). HSA and trypsin were selected for the initial validation. Both are available off the shelf, and binding of therapeutic proteins to HSA is a key determinant of pharmacokinetics. Therefore, single-domain antibodies targeting HSA may provide a modular tool for enhancing the half-life of biologics (*16*). Conversely, trypsin offers the opportunity of testing the design strategy on poorly accessible convex epitopes harbouring an active site. The RBD of SARS-CoV-2 exemplifies the power of targeting specific epitopes, as binding to regions overlapping with, or close to the ACE2 receptor binding site, whilst avoiding glycosylation sites, is known to yield neutralising antibody candidates, which would sterically hinder virus binding to the human cell receptor (*17*). In this case, we used as starting point for the design the first-released cryo-EM model of the SARS-CoV-2 spike protein in the prefusion conformation (*18*) (PDB ID 6VSB). The reason for this choice was to assess how the design strategy performs with a lower resolution structure used as input. Specifically, we ran the design on the surface of the up RBD around the ACE2-binding region, which has some regions of very low resolution (∼6-8 Å) (*18*), and several missing residues in the model (**Fig. 4A**).

**Table 1.**
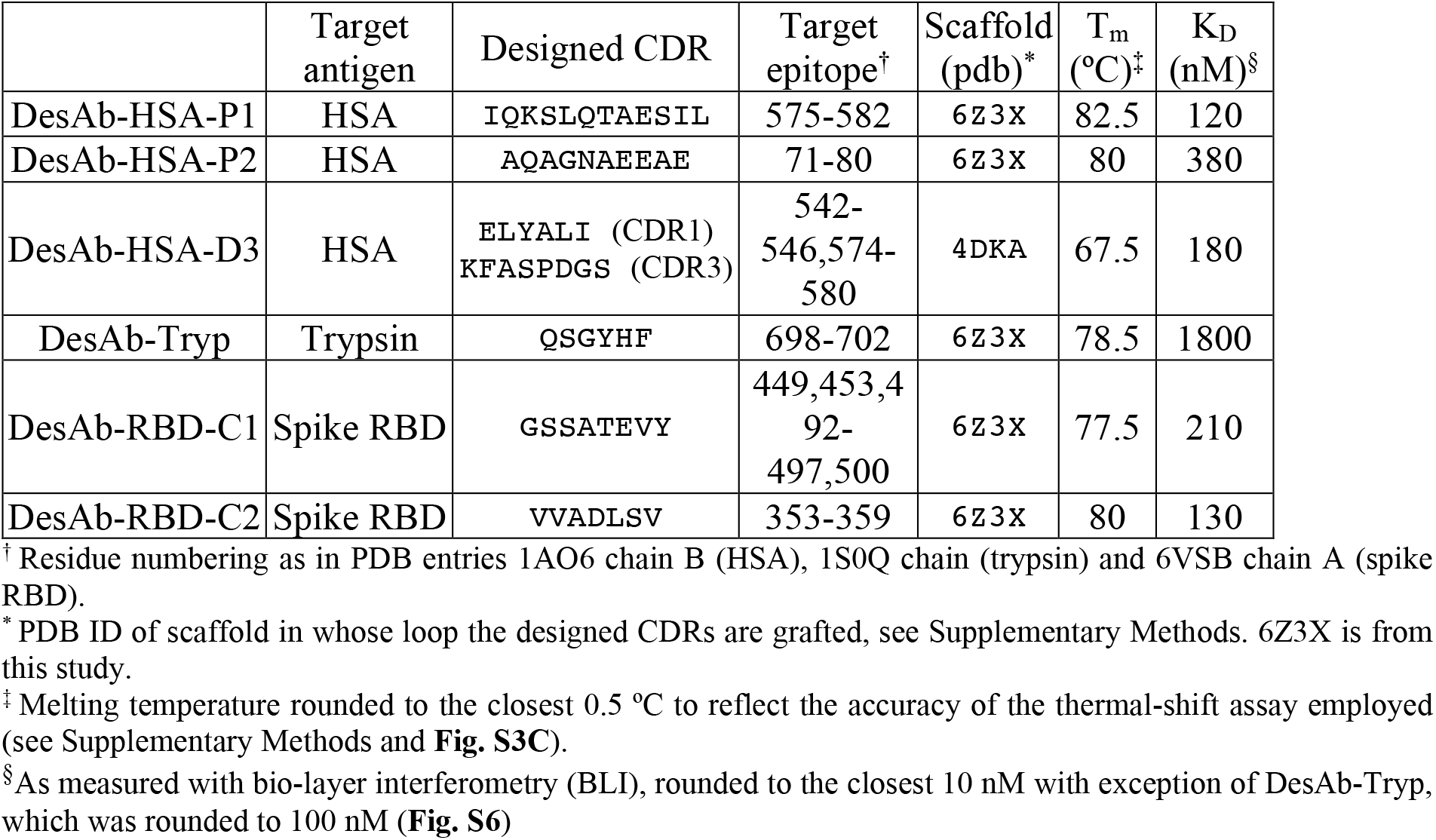
Designed single-domain antibodies (DesAbs) employed in this study.

All designed single-domain antibodies expressed well in *E. coli*, were obtained to high purity, and showed circular dichroism (CD) spectra fully compatible with a well-folded VH domain (**Fig. S2**, see Supplementary Methods). All designs were highly stable, with a melting temperature at par or better than that of immune-system-derived nanobodies (*19*) (**Table 1** and **Fig. S2C**).

Two out of the three anti-HSA single-domain antibodies, DesAb-HSA-P1 and DesAb-HSA-P2 (**Table 1** and **Fig. 2D**), consisted in designed CDR motifs grafted in place of the CDR3 of a previously characterised single-domain antibody scaffold highly amenable to CDR3 substitutions (*20, 21*) (**Table S1**). The third design, DesAb-HSA-D3, was made by structurally matching two separate CDR-like candidates onto two CDR-loops of a nanobody scaffold identified as highly compatible with these two binding motifs (**Fig. 2D**, see Supplementary Methods). The first strategy provides the opportunity to test the de novo CDR design procedure by minimising possible complications arising from the grafting, while the second is a more complex approach that allows to design multiple CDR loops onto a scaffold structurally matched to the epitope.

Binding to HSA was measured in solution with micro-scale thermophoresis (MST), which yielded KD values ranging from 140 to 800 nM (**Fig. 3A-C,E**), while a control single-domain antibody showed extremely weak signal in this assay (**Fig. S4A**). To put this in context, a nanobody isolated with yeast-display from a state-of-the-art naïve library, called Nb.B201, was recently reported to bind HSA with a KD of 430 nM (*22*), which is in the same range as those of our de novo designs.

**Figure 3.**
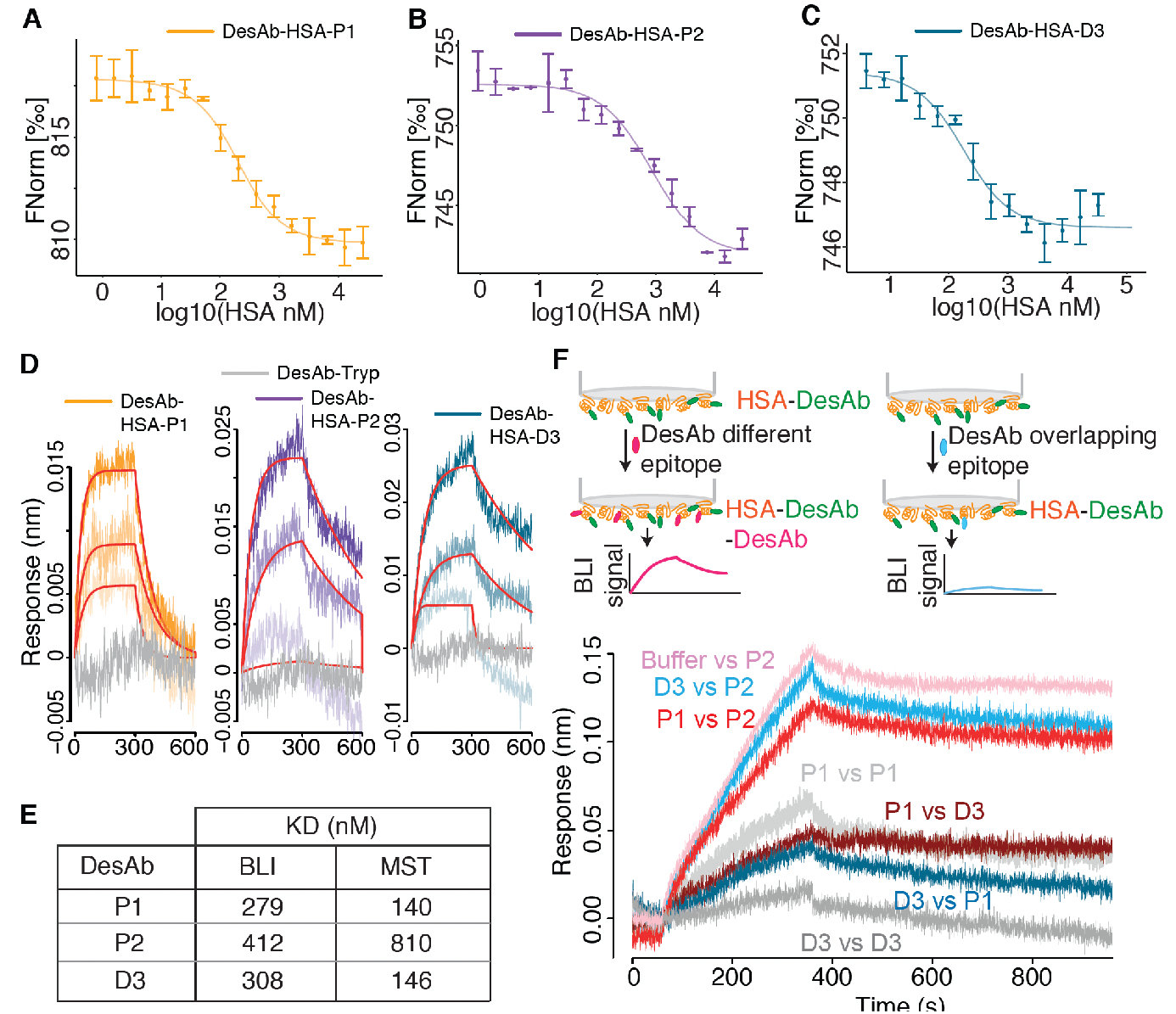
The anti-HSA DesAbs bind their target and compete for binding to overlapping epitopes. (**A**-**C**) Micro-scale thermophoresis (MST) of fluorescently labelled DesAbs (70 nM) in the presence of increasing concentrations of HSA (x-axis). Data points are mean and standard deviations of three replicates, data were fitted with a single-site binding model. **(D)** Biolayer interferometry (BLI) binding traces (association and dissociation) obtained with APS-sensors loaded with HSA. Association was monitored in wells containing 1 μM, 0.5 μM, and 0.25μM DesAbs. 1 μM DesAb-Tryp is shown in grey and was employed as control for non-specific binding to the sensor. (**E**) Table with the dissociation constants (KD) obtained for the three DesAbs by fitting the BLI and MST data. (**F**) Binding competition experiment at the BLI. APS-sensors were loaded with HSA, and then dipped in wells containing 5 μM of a first DesAb *X1* (see Supplementary Methods), then moved in buffer wells for one minute, and then into wells containing 5 μM of a second DesAb *X2*, and finally back to buffer wells. Curves are labelled with “*X1 vs X2”* to identify the anti-HSA DesAbs employed. The plot shows the last three steps, and reference sensors monitoring the background dissociation of *X1* during these steps were subtracted from the traces shown here. The traces *P1 vs P1* and *D3 vs D3* were taken as positive controls for the competition, and the small signal observed is due to the facts that not all epitopes are occupied by the first DesAb (*X1*), and that this is dissociating in the background. The trace *Buffer vs P2* was taken as a negative control for the competition.

To confirm the binding, we also carried out bio-layer interferometry (BLI) with immobilised HSA, obtaining KD values compatible with those measured in solution (**Fig. 3D,E**). Importantly, the trypsin-targeting DesAb-Tryp employed as a negative control gave no binding signal for HSA in this assay (**Fig. 3D**), while the yeast-display-derived anti-HSA nanobody Nb.B201, employed as a positive control, yielded a KD compatible with that reported in the literature (*22*) (**Fig. S4B**). DesAb-Tryp has the same sequence as DesAb-HSA-P1 and P2, except for the designed CDR motif grafted in the CDR3 loop (**Table S1**), and therefore represents a particularly suitable negative control to confirm that the observed binding is indeed coming from the grafted designed motif. Besides, DesAb-Tryp was able to bind its intended target trypsin, while DesAb-HSA-P1 and P2 showed no binding signal and were likely partly digested by the protease during the binding assay (**Fig. S5**).

The crystal structure of DesAb-HSA-P1 in the unbound form further confirms the correct folding of the VH domain. This structure also reveals the dynamic nature of the CDR3 loop, which harbours the designed motif, as the electron density is missing for most of this region (**Fig. S3**). A highly dynamic CDR3 loop was expected for this scaffold. For example, two of the four identical chains comprising the asymmetric unit of the structure of the original single-domain scaffold (PDB ID 3B9V) also have unassigned coordinates in their CDR3, even if the loop here is 8 residues shorter than that of DesAb-HSA-P1. The highly dynamic nature of this loop likely stems from the lack of strong CDR3-framework contacts, which is why folding and stability of this scaffold have been shown to be insensitive to mutations in its CDR3 loop by several studies (*20, 23*–*26*). Indeed, we selected this scaffold precisely because it can harbour virtually any sequence in its CDR3 without drastic consequences on its stability. However, the dynamic nature of the loop harbouring the designed motifs likely also explains why we were unable to obtain a crystal structure of DesAb-HSA-P1 bound to HSA. We speculate that such dynamic loop, even when bound to the antigen, retains enough hinge flexibility to embody the resulting complex with a degree of dynamics unsuitable for structural determination.

In the absence of an atomic-level structural characterisation of the designed interaction, we resorted to epitope binning through competition experiments. BLI competition experiments show that DesAb-HSA-P1 and DesAb-HSA-D3 compete with each other for binding to HSA, as the binding of one is hindered by the presence of the other antigen-bound DesAb (**Fig. 3F**). Conversely, DesAb-HSA-P2 does not compete with the other two, as its binding is not affected by the presence or absence of other antigen-bound DesAbs (**Fig. 3F**). This competition behaviour is fully compatible with the rational design, as DesAb-HSA-D3 and DesAb-HSA-P1 were designed to target partly overlapping epitopes, while DesAb-HSA-P2 targets a different epitope on the opposite side of the antigen (**Fig. 2D**).

Like the HSA-targeting DesAbs, the two designs made to target the RBD of the spike protein showed binding in the nanomolar range. We first tested the binding in solution to the full trimeric spike protein using MST (**Fig. 4B**, see Supplementary Methods). Both RBD-targeting DesAbs showed binding to the spike protein, while the HSA-targeting DesAb-HSA-P2 employed as a negative control gave no signal in the assay (**Fig. 4C**), confirming that the observed binding indeed comes from the designed CDR3 motif. Fitting the binding curves with a 1:1 binding model reveals apparent KD values of 150 and 580 nM, respectively for DesAb-RBD-C1 and DesAb-RBD-C2. As the spike protein is a trimer, a 3:1 binding model would have been theoretically more suitable. However, while three distinct drops may be discernible in the binding curve of DesAb-RBD-C1, these are largely absent from that of DesAb-RBD-C2, and in both cases the error bars are too large for a reliable 3:1 fit. To confirm the binding, we carried out a BLI assay with immobilised natively glycosylated RBD, which yielded KD values of 210 and 130 nM for DesAb-RBD-C1 and DesAb-RBD-C2, respectively (**Fig. 4D-E**). Conversely, these two anti-RBD antibodies showed no binding signal for immobilised HSA employed as a negative control and as a blocker in the assay (**Fig. S4C**, see Supplementary Methods). It is worth noticing that the lower apparent affinity of DesAb-RBD-C2 for the full spike, together with the absence of a three-step transition in its MST binding curve, is compatible with DesAb-RBD-C2 having a more sideways epitope (**Fig. 4A**), which may be poorly accessible in the down RBD conformation of the full spike (*18*). Finally, both anti-RBD DesAbs were able to compete with the binding of the human ACE2 receptor to the viral RBD, which suggests that affinity-matured versions of these DesAbs may have neutralizing potential (**Fig. 4F**).

**Figure 4.**
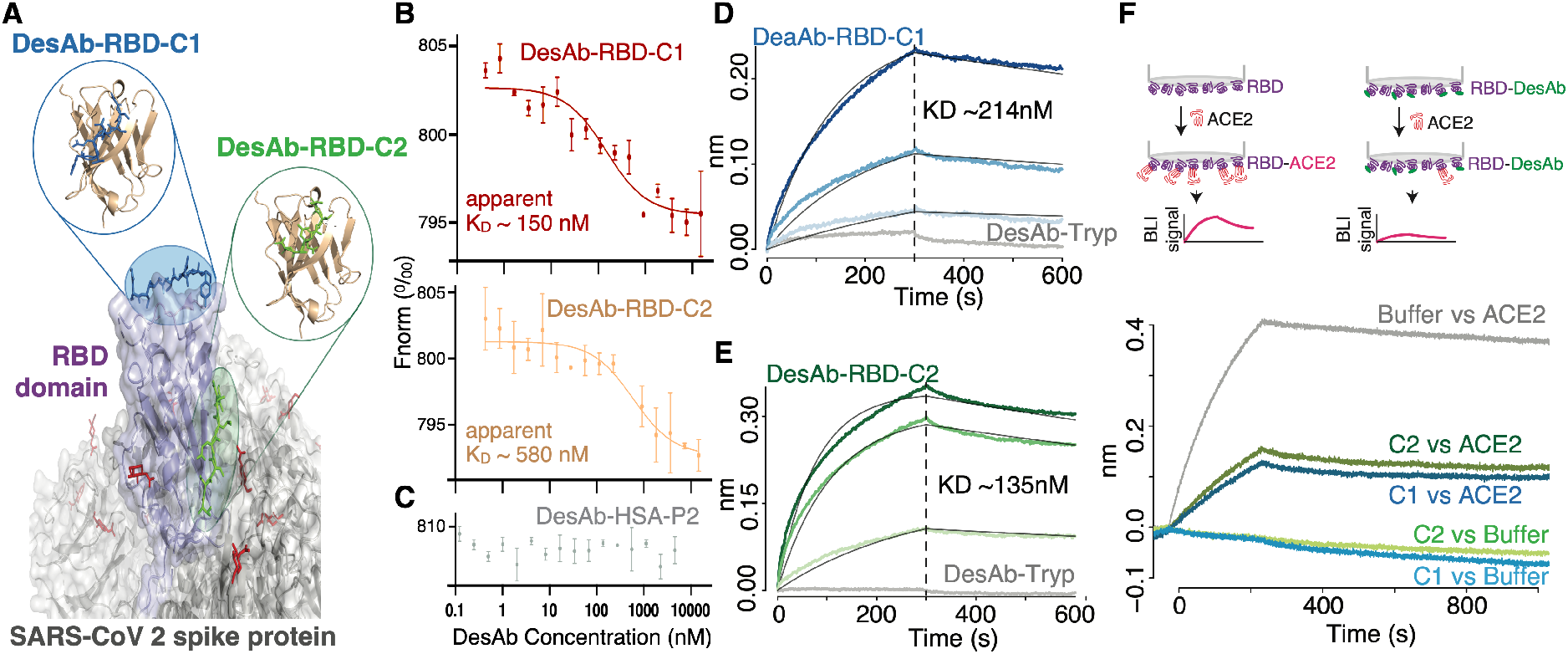
The anti-RBD DesAbs bind their target and compete with human ACE2. (**A**) the RBD is shown in purple, and the rest of the SARS-CoV-2 spike protein in grey with glycans in red. The designed CDRs targeting the RBD are in blue and green respectively for DesAb-RBD-C1 and C2, and corresponding structural models of the single-domain antibodies are represented in the bubbles. (**B, C**) Micro-scale thermophoresis of Alexa647-labelled trimeric SARS-CoV-2 spike protein (8 nM) in the presence of different concentrations of DesAb-RBD-C1 (red), DesAb-RBD-C2 (yellow), and (**C**) DesAb-HSA-P2 (gray) used as a negative control. KD values reported are obtained by fitting a 1:1 binding model, but they are only apparent KD ‘s as each spike has three RBD domains that can in principle bind independently to the DesAbs. (**D, E**) BLI binding traces (association and dissociation) obtained with APS-sensors loaded with RBD and blocked with HSA. Grey traces are obtained with 4 μM of DesAb-Tryp used as a negative control to probe non-specific binding to the sensors. (**B**) 4 μM, 2 μM and 1 μM of DesAb-RBD-C1 (from darker to lighter blue, KD= 214 ± 4 nM). (**C**) 4 μM, 2.5 μM and 1 μM of DesAb-RBD-C2 (from darker to lighter green, KD= 135 ± 2 nM). Data were fitted globally to estimate the reported KD values. (**F**) Binding competition experiment at the BLI. Sensors were loaded like in (D, E) and dipped in wells with 5 μM DesAb-RBD-C1 (blues) or DesAb-RBD-C2 (greens) or buffer (grey), then in wells containing ACE2 or buffer controls (see legend), and finally back to buffer. The plot reports the last two steps, showing that the binding of ACE2 is substantially reduced by the presence of either DesAb-RBD-C1 or DesAb-RBD-C2 bound to the RBD.

## Applicability of the design strategy

Having established that our computational method can yield stable single-domain antibodies that bind their intended targets with KD values down to the nanomolar range, we asked how readily and generally applicable the design strategy is. Given the fragment-based combinatorial nature of our method, we first asked what are the chances that suitable CDR-like fragments can be designed to target a given epitope, i.e. how typical it is for an epitope to have appropriate matching fragments in the *AbAg database*. To address this question, we run our design pipeline on the whole surface of all experimental target structures from the Critical Assessment of Techniques for Protein Structure Prediction competition (CASP14) (*27*). The target structures of the CASP assessments are selected ensuring that they represent a diverse sample of native folds, characterized by different sequences, secondary structures, and overall shape (*28*). Therefore, these structures also constitute a particularly suitable test set to explore the applicability of our design strategy. Having obtained all possible designed CDRs for each structure, we computed the solvent accessible surface area (SASA) of the structure in the presence and absence of bound designed CDR fragments, to reveal how much of the antigen surface is covered (see Supplementary Methods). Our results reveal that most of the surface of each antigen is typically targetable with our strategy, with a median surface coverage of 78% (**Fig. 5A**). Furthermore, for each epitope there are typically many candidate designed CDR loops to choose from, with a median density of 19 designed CDRs per nm^2^ of antigen surface (**Fig. 5B**). Taken together, these results reveal that, while some epitopes that cannot be targeted with our combinatorial strategy exist, most epitopes can be targeted by choosing between multiple different designed CDR candidates.

**Figure 5.**
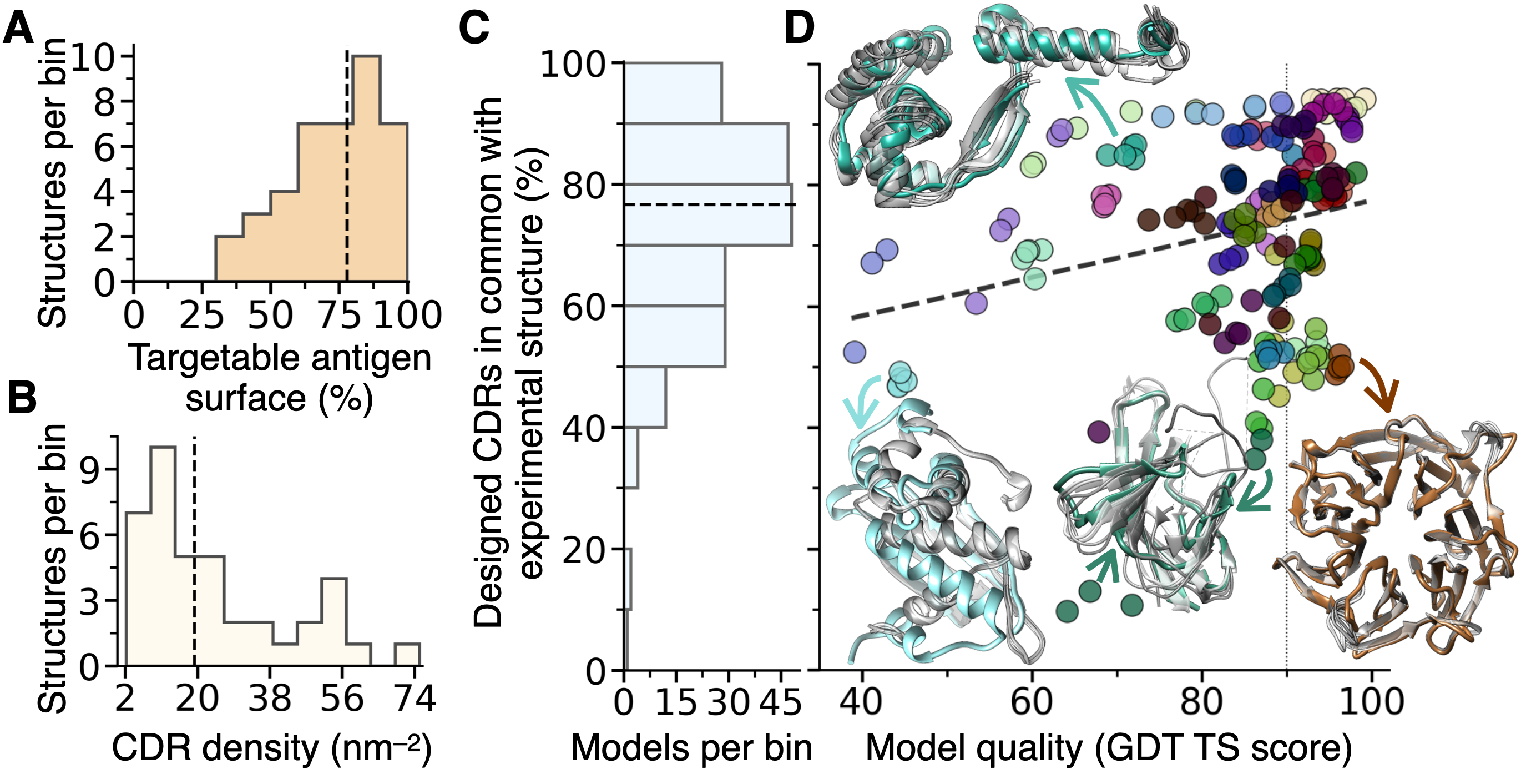
Applicability of the CDR-design procedure and performance on computationally predicted antigen models. (**A**) Histogram of the distribution of the percent of targetable surface area for each antigen (experimental structure from CASP14). Targetable surface is defined as the solvent-accessible surface area of the antigen made inaccessible by at least one designed CDR-like fragment. The dashed line is the median at 78%. (**B**) Histogram of the distribution of the CDR density for each antigen, expressed as the mean number of different designed CDRs per nm^2^ of antigen surface. The dashed line is the median at 19.2 CDRs per nm^2^. (**C,D**) Computationally predicted structural models generated by the AlphaFold2 algorithm within the CASP14 competition, as well as the corresponding experimentally determined structures, were used as input for the CDR design procedure. (**C**) Histogram of the distribution of the percent of designed CDRs obtained from each model that were identical to those obtained from the corresponding structure. The horizontal dashed line is the median of the distribution at 76.6%. (**D**) Scatter plot of the same CDR percent (y-axis) as a function of the global distance test total score (GDT TS, x-axis), as reported by the CASP14 competition, which is an indicator of the model accuracy. GDT works with the percentage of α-carbons that are found within certain cut-off distances of each other. A GDT of 100 means the modelled and experimental structure have all α-carbons within 1 Å of each other, and one above 90 (vertical dotted line) is typically regarded as a good solution of the folding prediction. The dashed trendline corresponds to a weak correlation (R^2^ = 0.06). Data points are coloured according to the target experimental structure of each model (see **Table S3** and **Fig. S6H** for a legend). Four example structures are drawn in the same colour as their model data points, which are pointed by the arrows. Their models are overlaid to the structures and shown in grey.

Having established that the vast majority of epitopes can be targeted with our design strategy, the most apparent bottleneck of the pipeline is the need for a structure to be used as input. As structural determination can be challenging for some antigens, this aspect could limit the applicability of the method, in particular in the cases of emerging diseases or of poorly investigated areas, where novel antibodies are often most needed. Recent advances in ab initio structure prediction are changing this scenario, as it is now possible to readily obtain rather accurate models of most protein structures of interest (*29, 30*). However, the accuracy of many methods of computational design, and in particular of those relying on energy functions that depend on interatomic distances, is known to rapidly deteriorate with lower-quality input models (*31*). Therefore, we next asked how applicable our method is on computationally modelled protein structures.

To test the dependence of our design method on the quality of the input structural model, we ran our CDR design procedure on all CASP14 models generated with AlphaFold2, which was the best-performing algorithm assessed (*27, 29*). By using all models deposited in CASP14 for each target structure, we make sure to include in our analysis also lower quality models that were not top-ranking in CASP (see Supplementary Methods). Our results reveal that most of the designed CDR-like fragments obtained by using each model as input are identical to those obtained using the corresponding experimentally determined structure (**Fig. 5A**). More specifically, the median number of designed CDRs in common between each model and its corresponding experimental structure, expressed as a percent of the total number of designed CDRs obtained for each model, is 77%, and only 20 (10%) of the 200 models analysed have less than 50% CDRs in common with their target structures (**Figs. 5A, S6** and **Table S3**). These results suggest that if one were to use an AlphaFold2 model as input for our antibody design pipeline, most typically about 75% of the generated CDRs would be identical to those that would be obtained from the corresponding crystal structure, and at least 50% would be identical in 90% of the cases. Besides, we only observe a very weak correlation (R^2^ = 0.06) between the percent of CDRs in common among model and structure, and the quality of the model itself as quantified by the global distance test total score (GDT, **Fig. 5B**). This result indicates that the aforementioned median number of designed CDRs in common among model and structure is not excessively determined by those very high-quality models (GDT ≥ 90) that are very similar to their target structure. Taken together, these results imply that the CDR-design procedure is expected to yield similar results when running on computer-predicted models or on experimental structures, and that these results do not strongly depend on the quality of the model used as input, at least within the quality range we explored (GDT > 40).

## Conclusions

We have described a fragment-based strategy for the rational design of antibodies targeting structured epitopes. We use protein fragments of at least four residues and typically longer in order to assemble designed CDRs in a combinatorial way. The idea behind this choice is that such fragments should be large enough to contain non-trivial sequence determinants of structure and interactions (*6, 20, 32*).

Our experimental results demonstrate that the design pipeline that we presented can successfully yield highly stable single-domain antibodies, which bind their intended targets with KD values down to the nanomolar range (**Table 1**). High nanomolar range binding (KD > 100 nM) was confirmed with two independent experimental techniques, one relying on equilibrium thermodynamics in solution (MST), and one on binding kinetics with a surface-immobilised ligand (BLI). We further verified, through various negative control experiments, that the DesAbs do not bind antigens that they are not intended for. Given that all DesAbs in this study, except for the two-loop design DesAb-HSA-D3, share the same framework sequence (**Table S1**), these experiments make us highly confident that the observed interaction is coming from the designed binding motif grafted in the CDR loop. Furthermore, we observed a binding-competition behaviour fully compatible with the location of the target epitopes on the antigen surface (**Figs. 3F** and **4F**). However, our failed attempts of obtaining a structure of the bound complex, together with the structure of DesAb-HSA-P1 (**Fig. S3**) suggest that these designed single-domain antibodies differ from immune-system-derived ones in their loop dynamics. Future work will be focussed on addressing this limitation, to enable the design of rigid DesAbs amenable to structural characterisation, which may even be applicable as crystallisation chaperones like natural nanobodies (*33*).

Importantly, we have been able to obtain DesAbs binding in the nanomolar range without the need of experimentally screening a large number of designs, but rather by pre-selecting in silico those designed CDRs that appeared most promising according to the metrics implemented, which include proxies for the predicted binding and sidechain complementarity, as well as predictions of solubility (*14*) (see Supplementary Methods).

The fragment-based combinatorial approach presented here does not involve approximations to calculate interaction free energies, and is also substantially faster than approaches based on the sampling of conformational and mutational space (*2*). An intrinsic limitation of this strategy, however, is that its applicability to epitopes of interest depends on the availability of suitable CDR-like fragments in the databases used. Nonetheless, the growing number of available protein structures in public databases makes the procedure generally applicable, as for most epitopes one obtains a number of candidate CDRs to choose from (**Figs. 5A,B** and **S1**).

Our results, which are obtained with a computer code that can run on standard laptops, demonstrate that it is becoming increasingly possible to design de novo antibodies binding to pre-selected epitopes of interest. We have exploited recent advances in protein-folding predictions and ab initio structural modelling to show that our design pipeline yields similar results when running on experimental structures or on computer-generated models, even when these do not reach atomistic accuracy. We envisage that, taken together, these advances in computational biotechnology will enable in the near future to obtain lead antibodies in a matter of days from the release of a pathogen genome, or from the identification of a novel disease-relevant target.

## Acknowledgements

We are grateful to Dr Paul Brear for useful advice on, and help with setting up protein crystallisation trials in Cambridge (Department of Biochemistry); to Dr Klara Kulenkampff for an aliquot of fluorescently labelled single-domain antibody (KK5 in Fig. S5); to Dr Faidon Brotzakis for useful discussions around the structural dynamics of the SARS-CoV-2 spike protein; to Matthias Schneider and Prof. Tuomas Knowles for useful discussion on, and help with biophysical measurements of protein interactions in solution; and to Dr Patrick Kuntz for sharing with us his dataset of nanobody melting temperatures (Fig. S2C).

P. S. is a Royal Society University Research Fellow (URF\R1\201461). Fondazione ARISLA (project TDP-43-STRUCT) and Italian Ministry of Research (PRIN 20207XLJB2) partially supported this work (S. R.). We acknowledge support from the Protein Interactions and Stability in Medicine and Genomics Challenge Programme (PRISM; NNF18OC0033950) funded by the Novo Nordisk Foundation.

