## Supplementary material for "Fragment-based computational design of antibodies targeting structured epitopes"

### COMPUTATIONAL METHODS

***Data collection.*** All protein structures in the Protein Data Bank [1] were downloaded from the rcsb.org website using a 90% sequence identity cut-off to reduce redundancy. Downloaded files were further cleaned by removing non-canonical amino acids and structures with no sidechain information. We refer to this dataset as the *PDB90 database*.

We further assembled a database of non-redundant complementarity determining regions (CDR), which we call *CDR database*. To create this dataset the Structural Antibody Database (SAbDab)[2] was downloaded, and the structures of all heavy and light CDR types (CDR-H1,2,3 and CDR-L1,2,3) according to the Chothia definition were extracted from the antibody structures and filtered for redundancy. A database of antibody-antigen structures, filtered for peptide or protein antigens only, was also obtained directly from the SAbDab website, and will be referred to as the *Antibody-Antigen database*. Finally, a database of complete structures of antibody Fv regions, comprising both variable heavy (VH) and variable light (VL) domains, as well as heavy-chain only antibodies (VHH) was retrieved from SAbDab and named *Ab database*.

***Generation of a database of antigen-cdr-like interactions.*** Each of the binding loop structures in the *CDR database* was used as query to look for structurally similar motifs in the *PDB90 database*. To achieve so, each template CDR loop of length *N* residues was fragmented using a sliding window approach with a range of [4, *N*] amino acids. Then, each of the generated fragments was matched against the whole *PDB90 database* using the MASTER program[3] (version 1.3.1) in order to find CDR-like structures. This structural search is based on the Kabsch algorithm[4], which employs root mean square deviation (RMSD) of the carbon alpha positions. For that, a RMSD cut-off of 0.4 Å was used for fragments of length 4, and increased by 0.05 Å for each additional residue (the maximum cut-off value was set to 1.0 Å). In this way, we obtained a database of CDR-like fragments whose backbone is found in a conformation compatible to that observed in at least one known antibody CDR, but with no constraints on sequence similarity with known CDRs.

Next, we sought to establish whether these CDR-like fragments had an antigen-like partner region in their native environments (i.e. in the structures where they have been identified). Here, we define antigen-like region any part of a protein structure within the *PDB90 database* comprising one or more fragments of at least 4 consecutive residues that is in contact with a CDR-like fragment. Two different definitions of contacting residues were used: first, those residues in the structure whose calculated solvent accessible surface area (SASA) increases upon removal of a CDR-like fragment; second, those fragments found within a distance of 10.5 Å between C$\alpha$ atom pairs from a CDR-like fragment. Therefore, as a final product two databases of antigen-like regions associated to the corresponding interacting CDR-like fragments were obtained, based on the two different residue-contact definitions. They will be individually referred to as the *AbAg-SASA* and *AbAg-CACA databases*, and collectively as the *AbAg database*.

***Identification of CDR-like fragments interacting with a structured epitope.*** Given the structure of an epitope of interest as input, the two *AbAg databases* can be searched to identify antigen-like regions structurally similar to those within the input epitope. In this way, the CDR-like fragments interacting with the identified antigen-like regions have the potential to also interact with the structure of the epitope used as a query, as long as these regions have a reasonable sequence similarity. In order to perform this search, the structure of the epitope is fragmented into smaller regions to increase the probability of identifying matching antigen-like regions in the databases. Two fragmentation modes are employed: the first one uses a sliding window approach to fragment contiguous peptides; window sizes are in the range [4,*N*], where *N* is the length of the input epitope, or the length of the epitope region under fragmentation in the case of input epitopes formed by multiple non-contiguous fragments. This fragmentation approach constitutes the "linear" mode. The other fragmentation mode takes each individual residue and calculates the closest *n* residues based on distances between the centre of mass of their side chains (**Fig. 1**). This is done with various *n* values in the range [4, *N*]. This fragmentation approach constitutes a conformational mode, as it can readily generate regions comprising non-contiguous polypeptide segments that are close with each other in the input structure of the epitope. All the generated fragmentations of the input epitope are used as queries to interrogate the *AbAg databases* using the MASTER program doing full backbone-to-backbone comparisons using the same RMSD metrics as those employed for the generation of the *AbAg databases*. To speed up the structural search, when using the linear fragmentation mode, the sequences of the generated epitope fragments are used as queries for a much faster blastp search[5] against the sequences of all antigen-like fragments within the *AbAg databases* (blast command: *blastp -query input_fragment_seauence.fasta -db AtAg_databases.fasta -qcov_hsp_perc 100.0 -matrix BLOSUM62 -task 'blastp-short' -word_size 2 -seg 'no' -evalue 20000 -ungapped -comp_based_stats F -max_target_seqs 60000 -outfmt 6 -out blast_hits.txt*). This strategy is used to restrict the search space of the MASTER program within the *AbAg databases* to only those antigen-like regions with a sequence identity meeting a user-selected threshold. When using the conformational fragmentation mode, sequence identity is checked during the *AbAg* structural search whenever a match is found. In both modes, whenever a matching antigen-like region meets both sequence identity and structure similarity criteria, the corresponding interacting CDR-like fragments are retrieved. While the sequence identity threshold is specified by the user, the RMSD threshold (in Angstroms) is given by the function $RMSDcutoff = 0.4 + n*0.033$, where *n* represents the number of residues in the epitope fragment used as query. The retrieved CDR-like structures are then rotated to match the orientation of the input epitope by superimposing the matching antigen-like region together with its interacting CDR-like fragment(s) to the input epitope. As the matching region is typically smaller than the full input epitope, steric clashes may occur between the identified CDR-like fragments and the rest of the epitope or of the antigen, in which case the CDR-like fragments are discarded. Otherwise, these are labelled as CDR-like candidates (**Fig. S1**).

***Optimization of the identified antigen-CDR-like interactions and raking of the hits.*** Each of the CDR-like candidates has a set of native interactions, which are defined as those interactions observed in the corresponding antigen-like region within the *PDB90 database* according to the SASA criterium of interaction described above. However, these interactions might not be fully conserved when the CDR-like candidate is paired with its corresponding epitope fragment, due to differences in amino acid sequence and side-chain orientation between the epitope fragment and the matching antigen-like region. If this is the case, the probability for the CDR-like candidate to interact with the epitope of interest might decrease. To address this issue, for each CDR-like candidate we run an optimization procedure on those residues that have different interactions with the input epitope than the corresponding native ones. For each of these CDR-like residues, the optimization starts by defining a local interacting structural motif. This motif comprises all epitope residues that are found interacting with the CDR-like residue under scrutiny according to the SASA criterium of interaction described above. Next, this local structural motif, which includes also the backbone atoms of the CDR-like residue itself, is used as a query to look for similar regions in the *PDB90 database* ($RMSDcutoff = 0.6 + n*0.025$, where $n$ is the number of residues). The aim is to find a matching region where the hit amino acid corresponding to the CDR-like residue under optimization has a backbone orientation very similar to the query, and therefore structurally compatible with the CDR-like candidate. Then, if those matching residues corresponding to the epitope residues have a sequence identity with the epitope higher than the current value, the sidechain of the CDR-like residue is replaced with that of the new hit, always avoiding hits that cause steric clashes or proline and cysteine residues that may respectively alter the CDR backbone conformation or later cause covalent dimerization of designed antibody candidates. This procedure is applied to all CDR-like candidates that need it, in order to maximise the number of native interactions. As multiple residue positions within each CDR-candidate may be optimized, and as each of them may have multiple optimization options, all possible combinations are generated. For example, a candidate with 3 optimisation options at position 1, and 2 options at position 4 will yield a total of 12 CDR-candidates.

All candidates are ranked according to their solubility, as computed by the CamSol method[6]. Furthermore, we also compute for each CDR-like candidate the number of native interactions, the number of shared interactions, and the number of interactions that are not shared, before and after optimization. Shared interactions are defined as interactions present in the original CDR-like/antigen pair found in the *PDB90 database* (native interactions) and that are also preserved in the optimized CDR-like bound to the epitope of interest. Based on these metrics, candidates with high number of shared interactions, low number of non-shared ones, and better solubility scores are regarded as the best ones. These scores and resulting rankings can be used to shorten the list of candidates and aid the selection of the most promising binding CDRs.

***Fragment assembly and CDR grafting.*** After optimization, the shortlisted CDR-like candidates are grafted in either full-length CDRs or directly full Fv antibody regions. At this stage, CDR-like candidates can also be combined with each other to obtain longer CDR candidates (**Fig. 1**). To do so, the first candidate of the shortlist is matched against the *Ab* or *CDR database* using MASTER, and the best match with no steric clashes between the epitope and the selected full CDR or complete Fv region is saved. Then, in order to combine together multiple CDR-like fragments in the same design, the same fragment is paired with all other fragments in the shortlist, and the pairs are matched against the *Ab* or *CDR databases* to see if both fragments could fit together different parts of the same CDR loop, or different CDR loops of the same Fv region. If any of the pairs of candidates is successfully matched, the result is taken to build triplets, and the matching process is repeated until no further match can be identified. After that, the process is repeated with the second candidate in the list, avoiding the already tested combinations. The iteration continues until all candidates and combinations are tested. Structural matching is done using Cα atoms comparisons with $RMSDcutoff = 0.4 + n*0.05$, where $n$ represents the total of residues in the query. This opens the opportunity of generating CDR loops comprising multiple CDR-like fragments, as well as antibodies with multiple candidates in different CDRs (**Fig. S1**).

This joining and grafting procedure may introduce new interactions between the CDRs in which the candidates where grafted and the epitope, and possibly also between the epitope and other parts of the Fv region. If that is the case, each new set of interacting residues on the antibody side is subjected to the optimization procedure described above in order to increase the chances of successful binding. Finally, the structure of the grafted candidates (either in CDRs or full Fv region) are produced as a final output.

***Generation of antibodies targeting human serum albumin, SARS-CoV-2 spike protein, and trypsin catalytic site.*** The described algorithm was applied to the entire surface of human serum albumin (HSA) (PDB ID 1AO6, chain B), to the antigen binding region of the receptor binding domain (RBD) of the SARS-CoV-2 spike protein (PDB ID 6VSB), and to a small region comprising the catalytic site of the trypsin protease (PDB ID 1S0Q). Both linear and conformational epitope fragmentation modes were employed, with 70% and 60% sequence identity thresholds used during the CDR-like candidate search, respectively. The search was constrained to fragments of length 4 to 13 amino acids. Both *AbAg databases* were used. The list of CDR candidates was shortened by selecting those whose number of shared interactions was greater than the number of non-shared interactions and corresponded to at least two thirds of the number of native interactions. For HSA, all shortlisted CDRs were then matched to full nanobody structures in order to find an amenable scaffold, and the top hit (based on the metrics describing the interactions, the solubility scores, and the quality of the grafting) was selected for experimental validation (DesAb-HSA-D3, consisting of 2 CDRs fragments matched to the CDR1 and the CDR3 of the VHH scaffold PDB 4DKA). Additionally, the two top hits from the shortlisted CDR candidates (DesAb-HSA-P1 and DesAb-HSA-P2) were taken to be grafted directly into the CDR3 of a stable VHH scaffold[7]. The latter strategy was also used for the Spike RBD designs (DesAb-RBD-C1 and DesAb-RBD-C2) and Trypsin active-site design (DesAb-Tryp).

**Analysis of the AlphaFold2 models and corresponding experimental structures within the CASP14 competition.**

Experimentally determined structures (targets) were downloaded from the CASP14 website, and specifically from https://predictioncenter.org/download_area/CASP14/targets/ (files therein were downloaded in January 2021 and last updated on 29 November 2020, e.g. casp14.targets.T-dom.public_11.29.2020.tar.gz). AlphaFold2 models were downloaded from the same website using the Table Browser feature, and by selecting ‘427 AlphaFold2’ at https://predictioncenter.org/casp14/results.cgi?view=tb-sel and 'all models' and 'all targets'. Selecting 'all models' instead of the default 'model 1' is important as it enables to assess multiple models for each experimental target, including those that were not top ranking and hence have lower quality. This table also contained all the model-quality metrics as calculated by the authors of the CASP14 competition, such as the root-mean-square deviation (RMSD) and the GDT_TS score (Global Distance Test Total Score)[8,9] (**Table S3**).

Given that the experimental structures of some of the targets have not yet been released publicly, at the time of analysis coordinates were available for 31 different targets out of 63 expected from the table. As a consequence, we restricted our analysis to those 200 AlphaFold2 models that mapped on a target with available coordinates. These corresponded to 5 models per target, with the exception of 3 targets (T1024, T1030, T1038) that had a total of 15 models each, as for these targets two domains had been independently modelled (5 models per domain) and 5 additional complete models with both domains modelled together were generated, and of one target (T1050) that had a total of 20 models, as 3 domains had been independently modelled for it (again 5 models per domain plus 5 complete models). Furthermore, in a number of cases there were amino acid residues present in the AlphaFold2 models but not in the corresponding experimental structure (e.g. regions of missing electron density), or vice versa (residues not present in the model but present in the experimental structures). In these cases, we removed the extra residues before running the design calculations, so that these ran on models and corresponding structures containing exactly the same residues (**Table S3**, column 'Processed'). This was a necessary precaution as the presence or absence of stretches of residues can generate different designed CDRs when running the design calculations. All PDB files from structures and models were cleaned using the PDBcleaner tool available on our web server (www-cohsoftware.ch.cam.ac.uk) to remove HETATM and to grow any missing atom. It is worth noting that the authors of the CASP competition select their targets also ensuring that they represent a diverse sample of native folds characterized by different secondary structure contents and overall shape, thus making these structures a particularly suitable test-set to explore the generality of our antibody design strategy.

To obtain the results presented in the main text (**Fig. 5**), we ran our algorithm on the selected models and their corresponding experimental structures using a 70% sequence identity cutoff and the requested minimum length of the CDR-like fragments was set to 4 residues. We then calculated the SASA for the entire input structure as well as for the structure in complex with all the identified CDR-like candidates. Subtraction of these values indicates the "surface coverage" per input structure (**Table S3**).

In addition to the results presented in the main text, it is worth noting that the observation that data points corresponding to different models of the same structure tend to cluster together in **Fig. 5B** suggests that the nature of the antigen may play a bigger role in determining the robustness of the design procedure than the quality of the model itself. For example, the five lowest-ranking models, three of which are outlier in the distribution with less than 20% CDRs in common with their structure, are all for the same target (T1064 in **Table S3**, PDB ID 7jtl **Fig. 5A**, **B**). This is a viral protein with a long, disordered loop on one side and several missing residues, which are the main culprits for the large number of CDRs that are different among models and target. Furthermore, it is worth noting that (**Fig. S6**): (i) the overall number of designed CDRs is typically very similar between models and target (Pearson's R = 0.96), (ii) the fraction of designed CDRs that are obtained for the experimental structure and not for its models is typically small (median 17%), and (iii) the total number of designed CDRs for a model appears to correlate with the overall fraction of CDRs that would also be obtained from the experimental structure (R=0.51).

### EXPERIMENTAL METHODS

***Protein production and characterization.*** Genes encoding the anti-HSA single-domain antibody candidates (plus a C-terminal 7X His-Tag) were synthesized and cloned into an IPTG inducible vector (by Atum in vector PD444) including a leading OmpA sequence to enable translocation to the periplasm and ultimately facilitate intra-domain disulphide bond formation and the secretion of the product to the media. The anti-spike RBD and anti-trypsin designs were introduced via restriction free cloning into the CDR3 of the DesAb-HSA-P2 plasmid. For all designed single-domain antibodies, versions with a free C-terminal cysteine residue were created using site directed mutagenesis. This cysteine was inserted as part of an Asp-Cys-Glu motif right before the start of the C-terminal HisTag. For the anti-spike RBD designs, versions with a π-clamp sequence

(FCPF)[10] followed by a TEV cleavage site right before the C-terminal 7X His-Tag were created by restriction free cloning.

Plasmids were transformed into *E. coli* Shuffle LysY strain to further facilitate the formation of the disulphide bond of the antibody. 0.5 L cultures of LB media were inoculated at initial 0.03 OD600 nm and were grown at 37 °C until reaching 0.8 OD600 nm, then induced with IPTG 500 µM. Overnight expression was carried out at 30 °C. Cellular pellet was discarded after centrifugation, and the supernatant was filtered using a 0.45 µm filter to remove remaining cell debris. Supernatant was passed twice through a gravitational flow column packed with Ni Sepharose Excel IMAC resin (Cytiva, 17-3712-01) previously equilibrated in PBS pH 7.4. Then the column was washed with PBS pH 7.4, and with a gradient of imidazole in PBS (10 mM, 30 mM). The protein was then eluted at 200 mM imidazole. Fractions were analysed using SDS-PAGE and those with the most protein and highest purity were dialyzed against PBS to remove the imidazole. Purified proteins were diluted to 20 µM, aliquoted, flash frozen in liquid nitrogen, and stored at -80 °C. The positive control nanobody Nb.B201 was expressed in the same way but using temperature and timing described in the original work[11], and using the expression plasmid deposited in Addgene (pET26b_Nb.b201 Plasmid #131404).

***Antigens.*** Human serum albumin (HSA) was purchased from Sigma-Aldrich (A3782) as lyophilized powder, resuspended in PBS and further purified via gel filtration using a Superdex 200 SEC column prior to use in binding assays. Pancreatic bovine trypsin was purchased from Sigma-Aldrich (T1426). Recombinantly produced (from HEK293 cells) SARS-CoV-2 Spike Glycoprotein (S1) His-Tagged RBD was purchased from The Native Antigen Company (REC31849) and supplied to high purity in dry ice. SDS-PAGE analysis showed purity > 95% and a MW consistent with the fully glycosylated RBD (data from The Native Antigen Company). Human ACE2 (18-615) recombinant protein, used in the competition assay in **Fig. 4F**, was also purchased from The Native Antigen Company, where it was expressed in HEK293 cells with Sheep Fc-Tag (REC31876). Trimeric His-Tagged SARS-CoV 2 Spike Glycoprotein was purchased from The Native Antigen Company (REC31871-100). Protein and antibodies concentrations were determined by absorbance measurements at 280 nm using theoretical extinction coefficients calculated with Expasy ProtParam web server.

***Protein-thermal shift stability measurements.*** The melting temperature of the DesAbs was measured with a protein-thermal shift assay on a Bio-Rad CFX96 Touch qPCR machine using the ROX filter in white PCR plates. Samples were heated at 0.2 ºC/min from 25 to 95 ºC, and consisted in purified antibody in PBS at a final concentration of 8 μM, and of Sypro-orange dye in Protein Thermal Shift Buffer (Thermofisher 4461146) accounting for 25% of the final volume at a final concentration of 2x the recommended dilution. Sample volumes of 50 μL per well were used. The signal from the dye in the absence of proteins was subtracted from the sample signal before analysis. The melting temperature ($T_m$) was determined as the point of steepest derivative, the values reported in **Figure S2** are average and standard deviations over four replicates. In **Table 1** we chose to round the Tms to the closest 0.5 ºC as, while standard deviations across wells in the same plate are typically very small, inter-experiment variations tend to be slightly larger.

***Circular dichroism.*** Far-ultraviolet (UV) CD spectra of the DesAbs were recorded using a Chirascan Applied Photophysics spectropolarimeter equipped with a Peltier holder, using a 0.1-cm-pathlength quartz cuvette. Samples contained 6 μM protein in PBS. The far-UV CD spectra of all DesAbs were recorded from 200 to 250 nm at 25 °C, and the spectrum of the buffer was systematically subtracted from the spectra of all DesAbs to yield the plots in **Figure S2**.

***Maleimide labelling.*** In order to obtain conjugates of the design antibodies to Alexa-647 dye, the C-terminal cysteine variants of the design antibodies were incubated with 1 mM DTT for 10 minutes in order to reduce inter-DesAbs disulphide bond yielding covalent C-terminus-C-terminus dimers that may have formed during storage. DTT was then removed using Zeba desalting columns (ThermoFisher 89882), and samples were concentrated to 100 μM prior to incubation with Alexa-647-maleimide reagent (ThermoFisher A20347) for 1hr at room temperature. Free dye was removed using PD-10 desalting columns (Cytiva, 17-0851-01), and the labelling efficiency was assessed by absorbance measurements. Trimeric SARS-CoV 2 Spike protein was fluorescently labelled by incubating a 2.8μM protein solution with 50 molar equivalents of Alexa-647-NHS ester reagent (ThermoFisher A20006) in the dark during 2h at room temperature. Excess dye was removed by desalting three times with Zeba columns (ThermoFisher 89882) and labelling efficiency was determined by absorbance (estimated to be 13:1 dye to protein labelling ratio).

***Microscale thermophoresis (MST) binding affinity measurements.*** For the anti-HSA designs: starting from 30 µM HSA (150 µM for the KK5 control DesAb in **Fig. S4**), 16 samples of 1:1 serial dilutions were incubated with 70 nM Alexa-647 labelled antibody for 1 h at room temperature. Samples were prepared in 170 mM NaCl, 50 mM Tris-HCl, 10 mM MgCl2, pH 7.4 with 0.05% Tween-20. After incubation samples were run in triplicate in a Monolith NT.115 System (NanoTemper technologies) using 20% LED excitation power and 60% MST power, at 25 °C. For the anti-RBD designs: DesAb-RBD-C1 and DesAb-RBD-C2 (variants with the C-terminal π-clamp and TEV cleavage site) at 14.4µM and the anti-HSA control DesAb-HSA-P2 at 4µM were used as starting concentration for preparing 16 1:1 serial dilutions in PBS pH 7.4 with 0.05% Tween-20. They were incubated with a final concentration of 8nM Alexa-647-labelled trimeric SARS-CoV 2 Spike protein at room temperature for 1 hour. After incubation samples were run in triplicate in a Monolith NT.115 System (NanoTemper technologies) using 15% LED excitation power and 80% MST power, at 25 °C. All data was analysed and fitted using the Monolith System software assuming a 1:1 binding interaction.

***Biolayer interferometry (BLI) binding affinity measurements.*** BLI measurements were performed using an Octet-BLI K2 system (ForteBio). All assays were carried out in a black 96-well plate, 200 µL per well, all sensors were subjected to pre-hydration in the assay buffer for at least 15 min before usage. The assay plate was kept at 25 °C throughout the entire experiment. For consistency with the MST measurements, anti-HSA design binding assays were carried out in a buffer containing 170 mM NaCl, 50 mM Tris-HCl, 10 mM MgCl2, pH 7.4. First, two APS sensors (sample and reference) were pre-incubated in buffer for 15 minutes. Assay program consisted of a 150 s baseline in buffer, 300 s loading using 4 µM HSA, 300 s wash in buffer, 90 s baseline in buffer, 300 s association in 1µM, 500 nM, and 250 nM anti-HSA DesAbs for the sample sensor and buffer for the reference sensor, 300 s dissociation in buffer (**Fig. 2D**). As a control for non-specific binding to the sensors the same experiment was carried out with the DesAb-Tryp instead of the anti-HSA DesAbs (**Fig. 2D**). The positive-control Nb.B201 experiment was carried out in the same way but using 800nM and 400nM as analyte concentrations (**Fig. S4B**). Binding competition experiment of the anti-HSA designs was carried out in a similar way, in a buffer consisting for 1/3 of PBS pH 7.4 and for 2/3 of the aforementioned 50 mM Tris-HCl, 170 mM

NaCl, 10 mM MgCl2, pH 7.4. APS-sensors were loaded with HSA for 600 s, baseline for 300 s, and then dipped in wells containing 5 μM of a first DesAb X1 for 600 s, moved in buffer wells for 60 s, and then into wells containing 5 μM of a second DesAb X2 for 300 s, and finally back to buffer wells for 600 s to monitor dissociation. DesAbs X1 and X2 refer to different combination of the anti-HAS DesAbs as in the legend of **Fig. 2F**. Because trypsin could not be loaded effectively on APS sensor, the trypsin binding assay was carried out with Ni-Nta sensors using the same buffer composition as above. Sensors were loaded with 7.5 μM his-tagged DesAb-Tryp or control DesAb (either DesAb-HSA-P1 or DesAb-HSA-P2 as in **Fig. S5**) for 900 s. We found that loading these sensors to saturation was the only viable way to fully suppress the non-specific binding of trypsin to the nickel sensors, hence we systematically employed control DesAbs for all trypsin concentrations tested. These controls are identical to DesAb-Tryp except for the designed CDR3 (**Table 1**). Following loading, a baseline was taken for 180 s, then association and dissociation steps as in **Fig. S5**. Assays for DesAb-RBD designs were carried out in PBS with APS sensors, following the program: 120 s baseline in buffer, 90 s loading using 400 nM RBD, 300 s 4 μM HSA blocking, 12 0s baseline, 300 s association, 300 s dissociation.

Data with multiple DesAb concentrations were fitted globally with in-house python scripts, using Kon and Koff as global fitting parameters and Rmax as a local parameter (i.e. each DesAb concentration was allowed its own value of Rmax as these are probed by different BLI sensors). The $K_D$ was then derived as the ratio of Koff/Kon. Because of the shape of its dissociation curve, the positive control Nb.B201 experiment was fitted with a model that does not assume full dissociation at infinite time (**Fig. S4B**).

***Crystallization, data collection, data reduction, structure determination, refinement and final model analysis.*** DesAb-HSA-P1 was concentrated prior to the set-up of crystallization trials to a final concentration of 10 mg/ml. Crystals of DesAb-HSA-P1 were obtained with the vapour diffusion technique, in sitting drops, using equal volumes of the protein and of 0.1 M sodium cacodylate pH 6.5, 27% PEG 2000 MME as a precipitant solution. Diffraction data were collected on cryoprotected crystals (25% glycerol) at 100 K, at the I04 beamline of the Diamond Light Source using a 0.9795 Å wavelength. The collected dataset was processed with Dials[12] and Aimless[13] from the CCP4 suite[14]. The structure was solved by molecular replacement with Phaser[15] using as a search model PDB ID 3B9V. The correct amino acids of the DesAb-HSA-P1 construct were built manually using COOT[16,17]. The initial model was refined alternating cycles of automatic refinement using Phenix[18] (version 1.17_3644) and manual model building in COOT. Data collection and refinement statistics are reported in **Table S2**. Analysis of molecular interfaces were performed using PISA[19].

***Accession codes.*** Coordinates of DesAb-HSA-P1 and structure factors were deposited to the Protein Data Bank with accession number 6Z3X.

| | |
|---|---|
| DesAb scaffold<br>(PDB ID 6Z3X) | MEVQLEESGGGLVQPGGSLRLSCAASGFNIKDTYIGWVRQAPGKGEEWVASIYPTSGYTRYADSVK<br>GRFTISADTSKNTAYLQMNSLRAEDTAVYYCAAGS**_DesignedCDR3_**EEFDYWGQGTLVTVSS<br>DHHHHHHH |
| DesAb-HSA-D3 | MEVQLQESGGGLVQAGGSLRLSCAASG_**ELYALI**_SMGWFRQAPGKEREFVAAISRNGDNTYYTDS<br>VKGRFTISRDNAKNTVELQMNSLKPEDTAVYYCAAD_**KFASPDGS**_VTIMTNEYDYWGQGTQVTVS<br>DHHHHHHH |

**Table S1**. Amino acid sequence of the single-domain antibodies employed in this study. In the sequence in the top row "_DesignedCDR3_" should be replaced with the designed CDR sequences reported in **Table 1** of the main text to obtain the sequence of each DesAb. The PDB ID 6Z3X is from this study. The 7x-His tag was used for purification. All DesAbs have a OmpA signal peptide, which is cleaved upon translocation to the periplasm yielding the sequences in the table.

| DesAb-HSA-P1 | |
|---|---|
| **Data collection** | |
| Wavelenght (Å) | 0.9795 |
| Space group | P2(1)2(1)2(1) |
| *Unit cell parameters* | |
| a, b, c (Å) | 40.72, 52.41, 99.18 |
| α, β, γ (°) | 90, 90, 90 |
| Wavelength (Å) | 0.9795 |
| Resolution (Å) | 46.34-1.74<br>(1.77-1.74) |
| Number of unique reflections | 45160 (2432) |
| $R_{merge}$ | 0.108 (0.776) |
| $R_{meas}$ | 0.121 (0.873) |
| $R_{pim}$ | 0.053 (0.396) |
| $<I/σ(I)>$ | 8.7 (1.5) |
| $CC^{1/2}$ | 0.998 (0.915) |
| Completeness (%) | 100 (99.4) |
| Multiplicity | 4.7 (4.7) |
| **Refinement** | |
| Resolution (Å) | 46.34-1.74 |
| Number of reflections | 42002 |
| $R_{work}/R_{free}$ (%) | 19.25 / 21.96 |
| *r.m.s. deviations* | |
| bond length (Å) | 0.010 |
| bond angles (°) | 1.049 |
| *Ramachandran plot* | |
| favored (%) | 99 |
| allowed (%) | 1 |
| outliers (%) | 0 |

**Table S2. Data collection and refinement statistics related to the structural determination of DesAb-HSA-P1.** Values in parentheses refer to the highest resolution shell.

**Table S3 (Included as separate tab-separated file). AlphaFold2 models and corresponding experimental structures from CAPS14 employed in this study.** This file can be opened with most spreadsheet editors including Microsoft Excel. Table with refernce to all model pdb files associated with their corresponding experimentally determined structures (Target pdb files). The model-quality scores in the columns 'GDT_TS', 'GDT_HA', 'RMS_ALL', and 'RMSD' were downloaded from the CASP14 website as calculated by the CASP authors. The column 'TP' (True Positives) reports the number of designed CDRs that could be obtained by using as input either the model or the experimental structure, 'FP' (False Positives) the number of designed CDRs obtained from the model but not from the stucture, and 'FN' (False Negatives) the number of designed CDRs obtained from the structure but not from the model. A small number of files (column 'Processed' set to 'Y') was processed by us to remove some residues that were present in the model and not in the structure or vice versa (see columns 'ResCoverage.of.original.model'and 'ResCoverage.of.original.structure'). All strutures and models were downlaoded from the CASP14 website (see Methods). 'precision', which is equal to TP/(TP+FP), is the fraction of CDRs obtained from the model that can also be obtained from the structure.

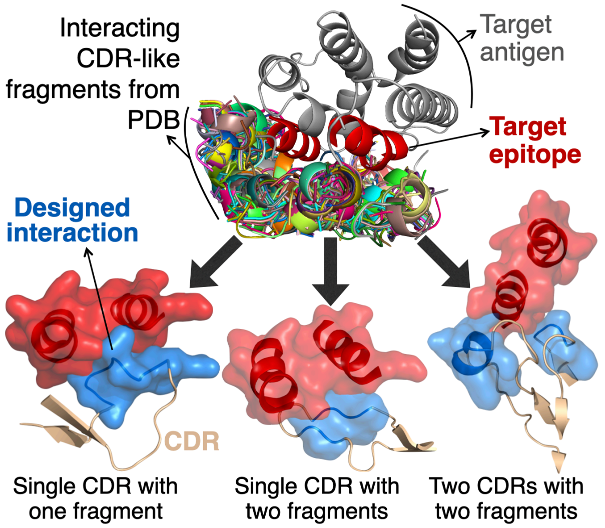


**Figure S1. Examples of designed CDRs to target conformational epitopes.** The target antigen is shown in grey at the top and the selected target epitope in red. Candidate interacting fragments (multiple colours) are selected as those fragments with a CDR-like backbone structure (e.g. backbone coordinates compatible with those of natural antibody CDRs) that are found in the Protein Data Bank (PDB) interacting with at least one fragment whose sequence and backbone structure match those of a corresponding fragment within the target epitope (see Methods). The lower panels, which are also reported in Fig. 2A of the main text, show three examples of how these CDR-like fragments can be combined in different ways to design interacting motifs (light blue), which are grafted onto structurally matched CDR loops (light brown) to generate lead antibodies predicted to bind to the target epitope.

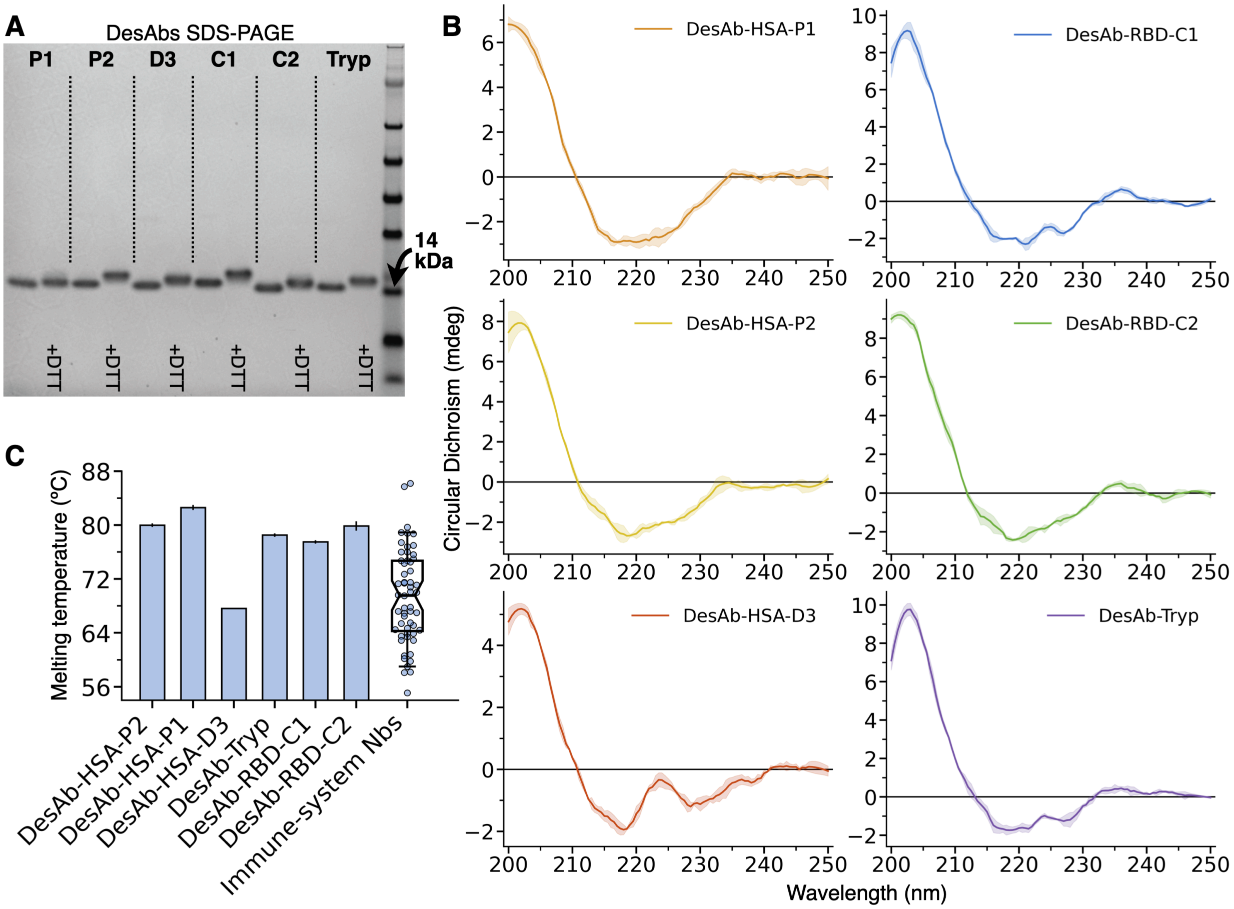


**Figure S2. Biophysical characterisation of the designed single-domain antibodies.** (**A**) SDS-PAGE of the six DesAbs designed in this study taken after purification (see legend), dashed vertical lines are guides for the eyes. The small difference in migration between samples without and with reducing agent (+DTT) indicates the correct formation of the intra-domain disulphide bond. (**B**) CD spectra of the six DesAbs (see legend) showing the expected minimum at 218 nm characteristic of β-domains. Profiles were smoothed using Savitsky-Golay filtering, and the shaded area is the standard deviation of three technical replicates. The spectrum of DesAb-HSA-D3 is substantially different from the others, as this design was done following a different strategy employing a different single-domain antibody scaffold for the grafting (see main text). (**C**) Bar plots with the melting temperatures of the six DesAbs measured with a protein thermal-shift assay (see Methods), compared with those of 68 nanobodies (swarm-plot to the right) isolated from camelids and reported in Ref. [20]. The box represents the first and third quartiles of the distribution of these 68 melting temperatures, whiskers represent the 1.5 interquartile range, and the bar at the centre the median.

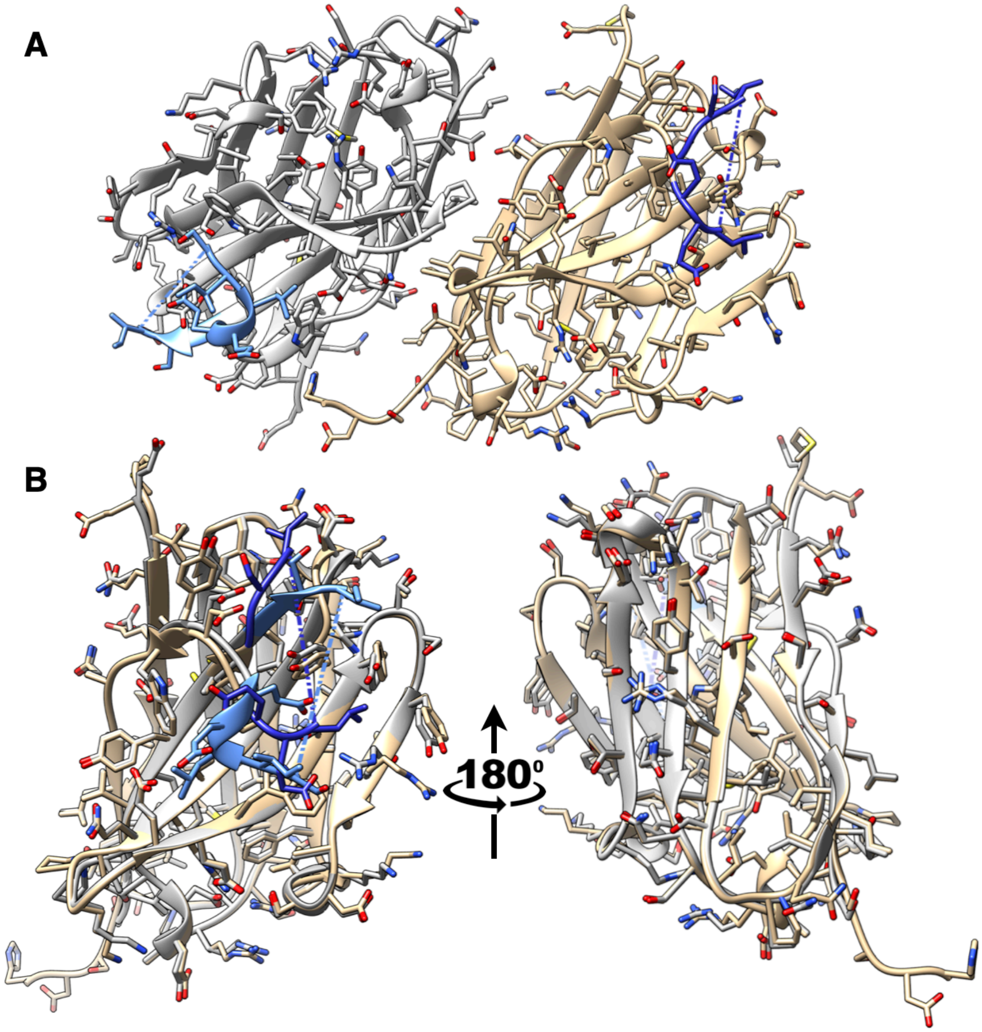


**Figure S3. Crystal structure of DesAb-HSA-P1.** (**A**) Asymmetric unit containing two single-domain antibodies, chain A is in grey and B in brown. The CDR3 loops that harbour the designed motifs are coloured in blue. Dashed lines denote a stretch of missing residues corresponding to the lower-case residues in the motif: `GSIqkslqtaeSILEE` (light blue in chain A) and `GSIqkslqtaesiLEE` (dark blue in chain B). (**B**) Structural superimposition of chain A and B from the asymmetric unit further reveals the dynamic nature of the CDR3 (blue), whose stems are found in substantially different conformations in the two single-domain antibodies. Coordinates have been deposited in the PDB with accession code 6Z3X.

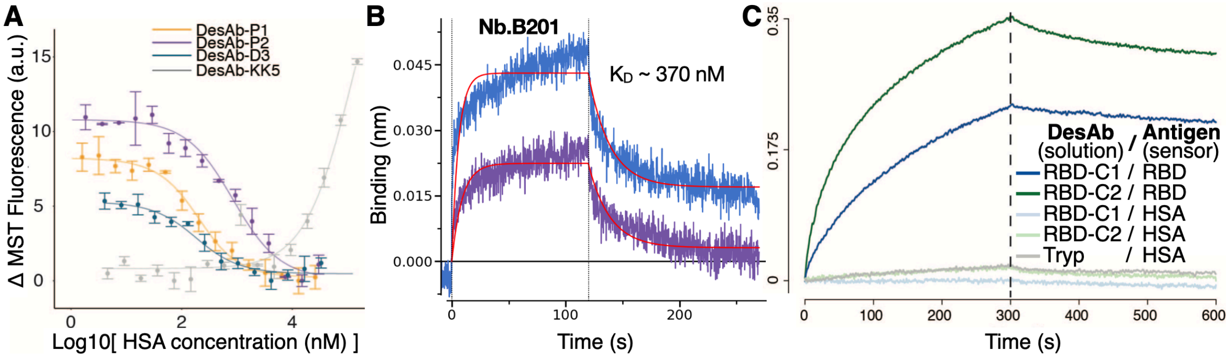


**Figure S4. Control experiments for the binding of the DesAbs to their targets.** (**A**) MST traces from **Fig. 3A-C** with the trace of an additional control single-domain antibody (KK5 in grey) targeting an unstructured epitope in an unrelated antigen (the human tau protein) and obtained as described in Ref. [7]. This single-domain antibody was fluorescently labelled with Alexa647 at an engineered solvent-exposed cysteine exactly like the other DesAbs (see Methods), and it is based on the same scaffold as DesAb-HSA-P1 and DesAb-HSA-P2 (VH-domain sequence identity respectively of 89% and 92%). For KK5 the signal deviates from flat at HSA concentrations >10 μM possibly because of molecular crowding and/or non-specific binding. Data for this control can be fitted with a $K_D \geq 110$ μM, which is only a lower bound as these data points do not reach a second plateau. (**B**) BLI experiment carried out exactly like those in **Fig. 3D** but using a positive control nanobody as analyte. The fitting model employed here does not assume full dissociation at infinite time and the KD obtained is in broad agreement with that reported in the literature (KD ~ 430 nM) [11]. (**C**) BLI experiment like in **Fig. 4D,E** using 4 μM of DesAbs in solution showing also binding traces obtained with different antigens on the sensors (see legend).

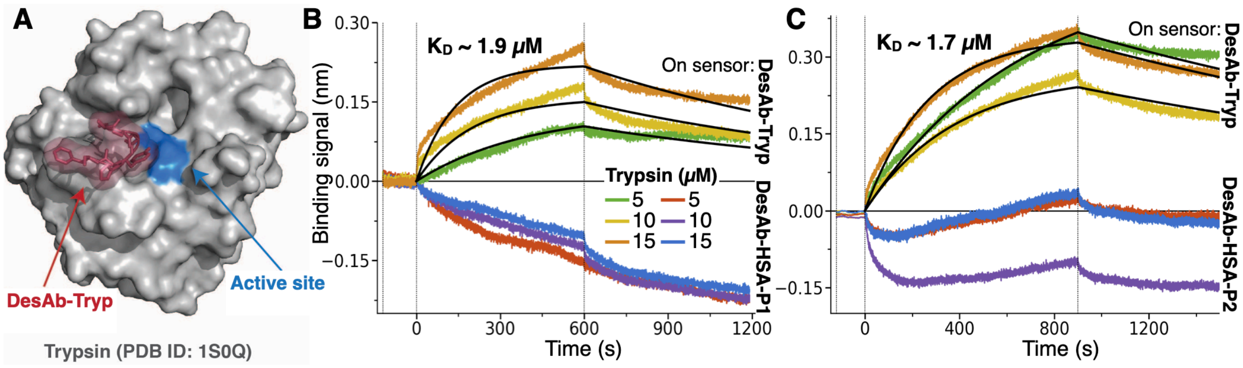


**Figure S5. Binding of DesAb-Tryp to trypsin.** (**A**) The structure of pancreatic bovine trypsin is shown in grey in surface representation (PDB ID 1S0Q), and the designed CDR fragment targeting an epitope within the active site (blue) is represented in dark red. This fragment is then grafted onto the CDR3 of DesAb-Tryp (**Table 1**). While running this design calculations, care was taken to exclude all known peptide inhibitors of trypsin or related proteases whose structure is available in the PDB. Indeed, because of their perfect complementarity with the active site of trypsin, such peptides figured as top-ranking, but testing them experimentally within a CDR loop would have defied the purpose of using our fragment-based combinatorial approach to design novel interactions. (**B**, **C**) BLI assays carried out with Ni-Nta sensors loaded with DesAb-Tryp (yellow, green and orange traces) or DesAb-HSA-P1 or DesAb-HSA-P2 (blue, red and purple traces, respectively in **B** or **C**). Panel **B** and **C** correspond to two independent experiments carried out in different days with a different choice of negative control DesAb. Shown are the end of the baseline phase, the association and dissociation phase separated by vertical dashed lines. Different Trypsin concentration are present in the association phase (see legend). Data from sensors loaded with DesAb-Tryp were fitted globally (see Methods), yielding a mean $K_D$ for trypsin of 1.8 μM. The decreasing signal observed for the negative control DesAbs is consistent with the trypsin protease digesting these $V_{HH}$s, which are immobilised on the sensor surface via their C-terminal His-tag. DesAb-HSA-P1is likely digested more, as, unlike DesAb-HSA-P2, it has a lysine in its designed CDR3 (**Table 1**).  DesAb-Tryp was not characterised further as we observed that, in solution, this was also digested by the trypsin protease.

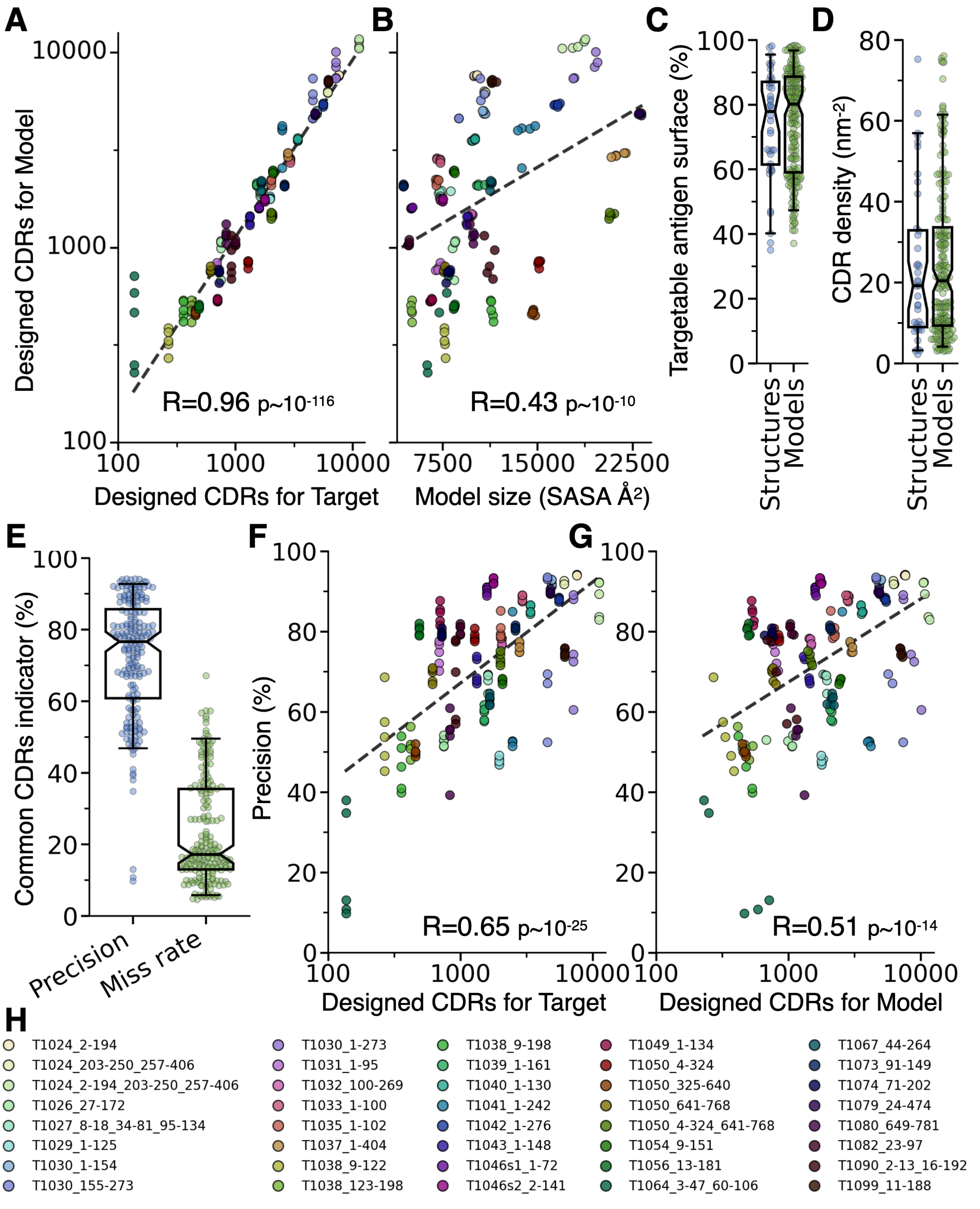


**Figure S6. Additional analysis of CDR-design on computationally predicted antigen structures.** (**A**) Scatter plot with the total number of designed CDRs that could be obtained when running the procedure using as input each structural model (y-axis) or its corresponding experimentally determined target structure (x-axis). (**B**) Total number of designed CDRs for each model as a function of the model size, expressed as its solvent-accessible surface area (SASA, x-axis). (**C**) Swarm plots of the percent of the input antigen surface that can be targeted with at least one designed CDR, and of the CDR density (**D**), calculated as the average number of designed CDRs per nanometre squared of the antigen surface. (**C**, **D**) Both quantities are calculated for all experimental structures (blue, same distributions as in **Fig. 5C** and **D** respectively) and all models (green). (**E**) Swarm plots of the percent of designed CDRs from each model that were identical to those from the corresponding target structure (Precision, same distribution as in **Fig. 5A**) and of the percent of designed CDRs from the target structure that were not found when running on each of its models (Miss rate). Boxes represent the first and third quartiles of the distribution, whiskers represent the 1.5 interquartile range, and the horizontal bar at the centre the median. (**F, G**) Scatter plots of the precision as a function of the total number of designed CDRs for each target structure (**F**) and for each model (**G**). In all scatter plots R is the Pearson coefficient of correlation and p its corresponding p-value. (**H**) Legend of the colour of the markers in this figure and in **Fig. 5**, which are coloured according to the identity of the target structure of each model (see **Table S3**). Labels are the CASP14 target ids underscore the residue-range that was modelled in each model.
